# Developmental regulation of kinetochore phosphorylation determines mitotic fidelity

**DOI:** 10.64898/2026.04.15.718713

**Authors:** Brian Galaviz Sarmiento, Duane A. Compton, Kristina M. Godek

**Affiliations:** Department of Biochemistry and Cell Biology, Geisel School of Medicine at Dartmouth, Hanover, NH, USA; Dartmouth Cancer Center, Geisel School of Medicine at Dartmouth, Lebanon, NH, USA

## Abstract

Accurate chromosome segregation relies on proper centromere and kinetochore formation and phospho-regulation. We previously demonstrated that a pluripotent state confers a low fidelity of chromosome segregation, however it is unknown how a pluripotent state impacts centromere and kinetochore function. Here, we demonstrate that both centromere and kinetochore structural organization and phosphorylation in mitosis are developmentally regulated. CENP-A, CENP-C, and HEC1 protein abundance is reduced at mitotic centromeres and kinetochores of human pluripotent stem cells (hPSCs) compared to isogenic somatic cells; however, elevating their levels does not improve chromosome segregation fidelity. Rather, we find that reduced phosphorylation of kinetochores is responsible for their low fidelity. HEC1 is hypophosphorylated at kinetochores of hPSCs compared to isogenic somatic cells at Cyclin B/Cdk1 and Aurora kinase phospho-sites. Inhibiting PP2A phosphatase activity or differentiation increases HEC1 phosphorylation at hPSC kinetochores decreasing chromosome segregation errors. Thus, mitotic fidelity in non-transformed human cells depends on the developmental regulation of the kinase and phosphatase networks controlling kinetochore phosphorylation.

**Summary:** Galaviz Sarmiento et al show that the developmental regulation of kinetochore phosphorylation governs mitotic fidelity. HEC1 is hypophosphorylated at kinetochores of hPSCs during mitosis contributing to their high rate of chromosome segregation errors. While differentiation increases HEC1 phosphorylation improving chromosome segregation fidelity.

## Introduction

Human pluripotent stem cells (hPSCs), including embryonic and induced, exhibit genomic instability during *in vitro* culturing and frequently become aneuploid with chromosome imbalances (Deng et al., 2023; Keller et al., 2019; Deckersberg et al., 2025). This is a major challenge for the efficacy and safe use of hPSC based treatments because aneuploidy impairs their ability to differentiate to other cell types and is a driver of tumorigenesis (Benvenisty et al., 2025). Aneuploidy results from chromosome missegregation during mitosis, and we previously showed that for hPSCs the primary cause of chromosome segregation errors is lagging chromosomes in anaphase. We also discovered that lagging chromosome rates correlate with developmental potential, increasing or decreasing with the gain or loss of pluripotency, respectively (Deng et al., 2023). These findings demonstrate that chromosome segregation fidelity is developmentally regulated and that a low fidelity is an inherent feature of a pluripotent state. Yet, how a pluripotent state affects the pathways responsible for ensuring accurate chromosome segregation is not understood.

Chromosome segregation relies on both the proper assembly of centromeres and kinetochores that form the binding sites for microtubule attachment to chromosomes and the establishment of correct bioriented kinetochore-microtubule (k-MT) attachments where sister chromatids are attached to microtubules from opposite spindle poles (Godek et al., 2014). We previously demonstrated that lagging chromosomes in hPSCs are caused by the persistence of erroneous merotelic k-MT attachments where a chromatid is simultaneously attached to microtubules from both spindle poles (Deng et al., 2023). This increases the probability of a chromosome missegregating and the generation of aneuploid daughter cells (Thompson and Compton, 2008; Ya et al., 2025; Deng et al., 2023; Cimini et al., 2001).

The frequency of erroneous merotelic attachments, and hence lagging chromosomes, partly depends on centromere and kinetochore size and protein abundance. Large kinetochores with increased levels of centromere/kinetochore proteins, including the histone H3 variant Centromere Protein A (CENP-A) that epigenetically determines the site of kinetochore assembly and the microtubule binding protein Highly Expressed in Cancer 1 (HEC1), are more prone to form merotelic attachments (Drpic et al., 2018). Conversely, reduction in CENP-A below ∼50% of wild-type centromeric levels also causes an increase in chromosome segregation errors, including lagging chromosomes (Fachinetti et al., 2013; Bodor et al., 2014). Thus, to achieve high mitotic fidelity there is an optimal range of protein abundance that supports centromere/kinetochore structural integrity and function. It was previously demonstrated that hPSCs have ∼50% decrease in centromere and kinetochore protein levels compared to somatic cells. This was postulated to effect chromosome segregation fidelity for hPSCs, but it was not functionally tested (Milagre et al., 2020).

In addition, the phospho-regulation of centromere and kinetochore proteins influences chromosome segregation error rates by facilitating the correction of merotelic k-MT attachments to proper bioriented attachments. Error correction relies on cycles of k-MT detachment and reattachment to promote bioriented attachment formation (Godek et al., 2014). A key player is HEC1 whose binding affinity for k-MTs is governed by electrostatic interactions through phosphorylation of at least ten N-terminal tail sites. Phosphorylation reduces HEC1 binding affinity for k-MTs to promote MT detachment and correction of improper attachments while hyperphosphorylation prevents any k-MT attachments from forming. Conversely, dephosphorylation increases HEC1 binding affinity for MTs. This facilitates k-MT attachment formation but suppresses detachment, resulting in a decrease in error correction and elevation in lagging chromosomes. The combined activities of kinases and phosphatases define the net phosphorylation status of each site to precisely “tune” total HEC1 phosphorylation levels and maintain k-MT attachment stability within a narrow dynamic range to promote faithful chromosome segregation (Kucharski et al., 2022; DeLuca et al., 2006, 2011; Zaytsev et al., 2014, 2015; Welburn et al., 2010). Here, we compare centromere and kinetochore protein abundance and phosphorylation levels between hPSCs and isogenic differentiated cells. Also, we functionally test if differences are responsible for the low chromosome segregation fidelity of hPSCs to shed light on the developmental regulation of mitosis.

## Results

### Mitotic centromere and kinetochore protein levels are low for hPSCs but sufficient for proper function

To investigate why a pluripotent state confers a high rate of lagging chromosomes, we compared mitotic centromere and kinetochore abundance of the histone H3 variant CENP-A, the constitutive centromere protein CENP-C that binds to CENP-A nucleosomes (Carroll et al., 2010), and the outer kinetochore protein HEC1 that binds to k-MTs for hPSCs and isogenic somatic cells. We used quantitative immunofluorescence microscopy to measure levels at individual centromeres or kinetochores from early prometaphase to anaphase comparing WTC-11 and GM human induced pluripotent stem cells to their respective isogenic primary somatic fibroblasts that were used for reprogramming. Also, we included H1 human embryonic stem cells. At all stages of mitosis, average CENP-A and CENP-C levels at centromeres were significantly reduced by ∼50% for WTC-11 and GM hPSCs compared to their respective isogenic fibroblasts. Average HEC1 levels at kinetochores of hPSCs were also significantly decreased but to a lesser extent. Similar trends were observed for H1 hPSCs compared to somatic fibroblasts (Figures 1A-C, S1A-C, S3A, and S4B). In agreement with our findings, centromere abundance of CENP-A and CENP-C in interphase cells and total nuclear HEC1 levels in mitotic cells were found to be lower for hPSCs vs. somatic cells (Milagre et al., 2020).

**Figure 1.**
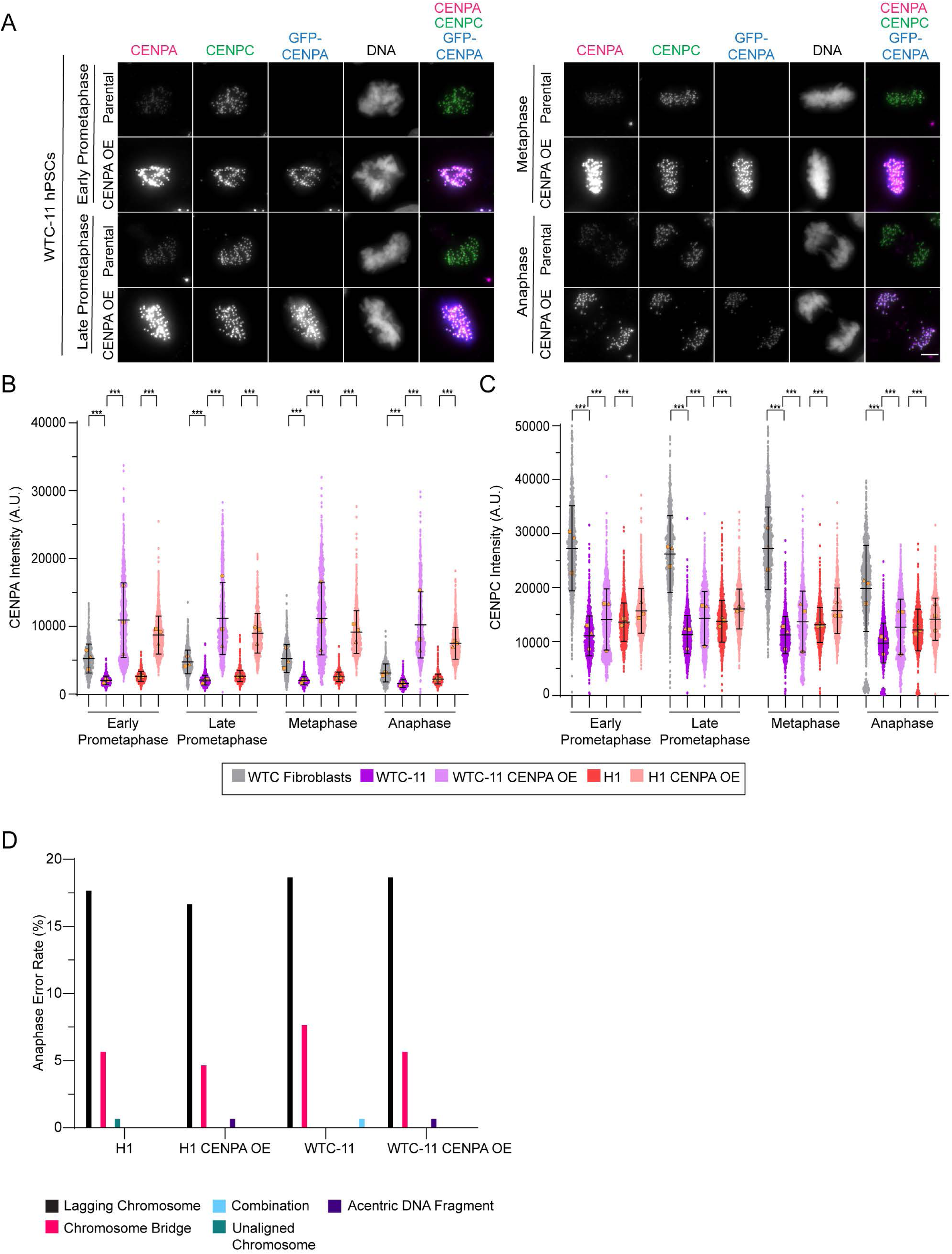
Mitotic centromere/kinetochore protein abundance is low for hPSCs but sufficient for proper function. (**A**) Representative images of mitotic parental WTC-11 and WTC-11 CENP-A-OE hPSCs from early prometaphase to anaphase. Shown is DNA (grayscale), CENP-A (magenta), CENP-C (green), and GFP (blue). The brightness and contrast for the CENP-A and CENP-C images are scaled equivalently between parental and CENP-A OE hPSCs for comparison. Scale bar 5 μm. (**B-C**) Quantification of total CENP-A (**B**) or CENP-C (**C**) kinetochore levels from early prometaphase to anaphase in WTC-11, WTC-11 CENP-A OE, H1, and H1 CENP-A OE hPSCs and WTC fibroblasts. Shown are individual kinetochore intensities and superimposed means of individual replicates (orange shapes). n > 900 total kinetochores from 20 cells per mitotic stage for each replicate with 15-25 kinetochores quantified per cell; mean ± SD. (**D**) Percentage of anaphase errors in H1, H1 CENP-A OE, WTC-11, and WTC-11 CENP-A OE hPSCs. Anaphase errors include lagging chromosomes, chromosome bridges, acentric DNA fragments, unaligned chromosomes, and a combination of errors. n > 300 anaphases. Three independent biological replicates (**B-D**); ***p < 0.001 using a one-way ANOVA with Holm-Sadak’s multiple comparison test (**B-C**) or two-tailed Fisher’s exact test with no significant differences to report (**D**).

A decrease in CENP-A below ∼50% of wild-type centromeric levels causes an increase in chromosome segregation errors for somatic cells (Bodor et al., 2014; Fachinetti et al., 2013), so we tested whether chromosome segregation fidelity would improve in hPSCs if centromeric CENP-A abundance was increased. We stably over-expressed GFP tagged CENP-A in WTC-11 and H1 hPSCs and confirmed that these cells maintained expression of pluripotency transcription factors (Figures S2A-D). For GFP-CENP-A hPSCs, average total centromeric CENP-A levels increased more than 4-fold above that of parental cells. This also elevated their total CENP-A levels above that of somatic centromeres (Figures 1A-B and S1C). CENP-C and HEC1 abundance also significantly increased in GFP-CENP-A hPSCs compared to parental cells. Interestingly, CENP-C levels were partially restored to somatic centromere levels in GFP-CENP-A hPSCs while at kinetochores HEC1 reached near similar levels (Figures 1A, 1C, S1C, and S4A-B). This indicates that CENP-A overexpression did not grossly alter centromere and kinetochore formation in hPSCs. We find that for hPSCs centromere and kinetochore protein abundance scales with the amount of CENP-A incorporated into chromatin, but scaling is not strictly proportional and varies for individual components.

We also measured interkinetochore distance (IKD) generated from microtubules pulling pairs of sister chromatids in opposite directions at metaphase to determine if CENP-A overexpression compromised centromere and kinetochore structural integrity and function. Increased IKD is indicative of centromere cohesion defects while decreased IKD results from an inability to form MT attachments (Sapkota et al., 2018; Zaytsev et al., 2014). Yet IKD remained similar between GFP-CENP-A and parental hPSCs (Figure S3B). Surprisingly, the increase in CENP-A centromeric chromatin resulting in CENP-C and HEC1 centromere/kinetochore abundance reaching near somatic levels, did not change chromosome segregation error rates, including lagging chromosomes, for hPSCs (Figure 1D). Collectively, these findings demonstrate the developmental regulation of mitotic centromere and kinetochore formation with respect to their stoichiometric organization. Moreover, although a pluripotent state reduces mitotic centromere and kinetochore protein abundance, protein levels are sufficient for proper structure and function indicating that alternative mechanisms are responsible for the low fidelity of chromosome segregation in hPSCs.

### HEC1 at kinetochores is hypophosphorylated at Cyclin B/Cdk1 and Aurora kinase sites for hPSCs

An alternative possibility is that a pluripotent state impacts the regulatory networks controlling mitotic fidelity. Our previous work showed that decreasing k-MT stability in hPSCs transiently reduces their lagging chromosome rate. This suggests that k-MT attachments are hyperstable in hPSCs relative to somatic cells, which decreases merotelic error correction causing their high rate of lagging chromosomes (Deng et al., 2023). The phospho-regulation of many centromere and kinetochore proteins influence k-MT attachment stability (Godek et al., 2014). Accordingly, we focused our attention on HEC1, a subunit of the NDC80 complex, because its N-terminal tail directly binds to k-MTs and phosphomimetic and non-phosphorylatable N-terminal tail mutants alone are sufficient to impact k-MT attachment stability and error correction (Kucharski et al., 2022; Zaytsev et al., 2014; DeLuca et al., 2006, 2011).

First, we analyzed phosphorylation of serine 44 in the HEC1 N-terminal tail using a phospho-specific antibody (DeLuca et al., 2011). We quantified the ratio of phosphorylated S44 to total HEC1 levels to control for the reduced abundance of HEC1 at kinetochores of hPSCs and for changes in HEC1 levels at kinetochores as mitosis progresses (Figures S3A and S4B). We found that HEC1 S44 was hypophosphorylated at kinetochores of WTC-11 and GM hPSCs compared to their isogenic fibroblasts, with ∼50% reduction across all stages of mitosis (Figures 2A-C). A similar trend was observed for H1 hPSCs compared to somatic cells (Figure 2C). In somatic and cancer cells, the absolute phospho-occupancy of HEC1 S44 changes from ∼50% to ∼25% between low and high affinity binding for microtubules (Kucharski et al., 2022). Thus, the reduction in S44 phosphorylation relative levels for hPSCs is within physiologic range to increase HEC1 binding affinity for k-MTs.

**Figure 2.**
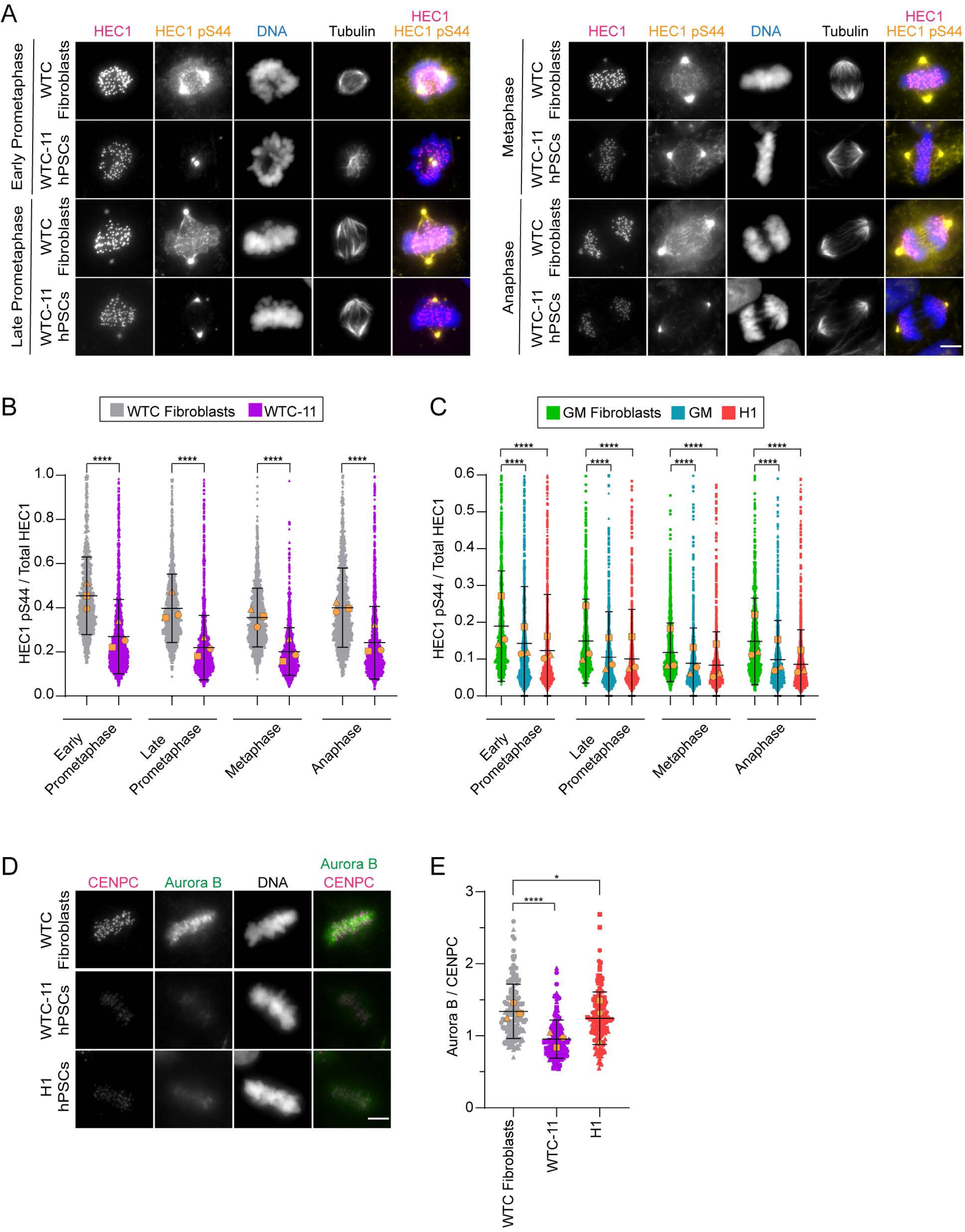
The HEC1 S44 Aurora kinase site is hypophosphorylated at mitotic kinetochores of hPSCs. (**A**) Representative images of mitotic WTC-11 hPSCs and isogenic WTC fibroblasts from early prometaphase to anaphase. Shown is DNA (blue), HEC1 (magenta), HEC1 phospho-S44 (orange), and tubulin (grayscale). The brightness and contrast for the HEC1 and HEC1 S44 images are scaled equivalently between WTC-11 hPSCs and WTC fibroblasts for comparison. Scale bar 5 μm. (**B**-**C**) Quantification of HEC1 S44 phosphorylation normalized to total HEC1 levels at kinetochores from early prometaphase to anaphase in WTC-11 and WTC fibroblasts (**B**) and GM hPSCs, GM fibroblasts, and H1 hPSCs (**C**). Shown are individual kinetochore intensities and superimposed means of individual replicates (orange shapes). n > 900 total kinetochores from 20 cells per mitotic stage for each replicate with 15-25 kinetochores quantified per cell; mean ± SD. (**D**) Representative images of metaphase WTC-11 and H1 hPSCs and WTC fibroblasts. Shown is DNA (grayscale), CENP-C (magenta), and Aurora B (green). The brightness and contrast for the CENP-C and Aurora B images are scaled equivalently between hPSCs and fibroblasts for comparison. Scale bar 5 μm. (**E**) Quantification of metaphase Aurora B levels in WTC-11 and H1 hPSCs and WTC fibroblasts. The maximum Aurora B intensity between sister kinetochore pairs was normalized to CENP-C. Shown are individual kinetochore pair intensities and superimposed means of individual replicates (orange shapes). n = 150 total sister kinetochore pairs from 10 cells for each replicate with 5 sister pairs quantified per cell; mean ± SD. Three independent biological replicates (**B**-**C** and **E**); *p < 0.05 or ****p < 0.0001 using an unpaired Welch’s t-test (**B**-**C** and **E**).

Next, we sought to determine if kinase activity was limiting at kinetochores of hPSCs relative to somatic cells explaining their decrease in HEC1 S44 phosphorylation. HEC1 S44 is proposed to be phosphorylated by Aurora B and/or Aurora A kinase with conflicting reports as to whether it is preferentially phosphorylated by Aurora B or Aurora A (Sobajima et al., 2023; DeLuca et al., 2011; Abe et al., 2016; DeLuca et al., 2017). Aurora A predominantly localizes to spindle poles, and not kinetochores, during mitosis to regulate spindle size. Amplified Aurora A activity causes an increase in spindle length (Sobajima et al., 2023), so we measured metaphase spindle size as a surrogate readout for Aurora A activity. Metaphase spindles were shorter in hPSCs compared to somatic cells consistent with reduced Aurora A activity (Figure S3C).

In contrast, Aurora B localizes to the cytoplasm, the inner centromere region between sister kinetochore pairs, and to kinetochores during prometaphase and metaphase (Broad et al., 2020). At kinetochores, tension generated from MT attachments pulling sister chromatids in opposite directions increases IKD and decreases Aurora B phosphorylation of HEC1 (Liu et al., 2009; Welburn et al., 2010; DeLuca et al., 2011; Kucharski et al., 2022). We measured IKD at pairs of metaphase sister chromatids and found that IKD was significantly shorter for WTC-11 and GM hPSCs compared to their isogenic fibroblasts. Likewise, IKD was shorter for H1 hPSCs compared to somatic cells (Figure S3B). This shows that elevated tension at kinetochores of hPSCs is not responsible for the decrease in HEC1 S44 phosphorylation. Also, we measured total Aurora B levels localized between CENP-C of sister kinetochore pairs at metaphase as previously described (Kucharski et al., 2022). We normalized Aurora B intensity to CENP-C to account for differences in overall kinetochore protein abundance. Aurora B kinase levels were reduced in WTC-11 and H1 hPSCs relative to WTC fibroblasts (Figures 2D-E); however, there was not a consistent difference. H1 hPSCs had only a marginal decrease in Aurora B abundance compared to WTC fibroblasts while WTC-11 hPSCs had a greater decrease (Figure 2E). These findings indicate that reduced levels and activities of Aurora kinases contribute to the diminished phosphorylation of HEC1 S44 for hPSCs.

We were curious as to whether HEC1 phosphorylation differences between hPSCs and somatic cells extended to other non-Aurora kinase N-terminal tail phospho-sites, so we measured the levels of phosphorylated HEC1 at threonine 31 using a phospho-specific antibody. HEC1 T31 is phosphorylated by the Cyclin B/cyclin dependent kinase 1 (Cdk1) complex to facilitate the correction of erroneously attached k-MTs (Kucharski et al., 2022; Kettenbach et al., 2011). Similar to the S44 site, HEC1 T31 was hypophosphorylated with ∼50% reduction at kinetochores of hPSCs compared to somatic cells from early prometaphase to anaphase (Figures 3A-C). Also, consistent with chromosome segregation error rates remaining elevated, HEC1 T31 was hypophosphorylated in GFP-CENP-A expressing hPSCs relative to somatic cells (Figures S4A-B). Moreover, the reduction in T31 phosphorylation relative levels for hPSCs is within physiologic range to increase HEC1 binding affinity for k-MTs (Kucharski et al., 2022).

**Figure 3.**
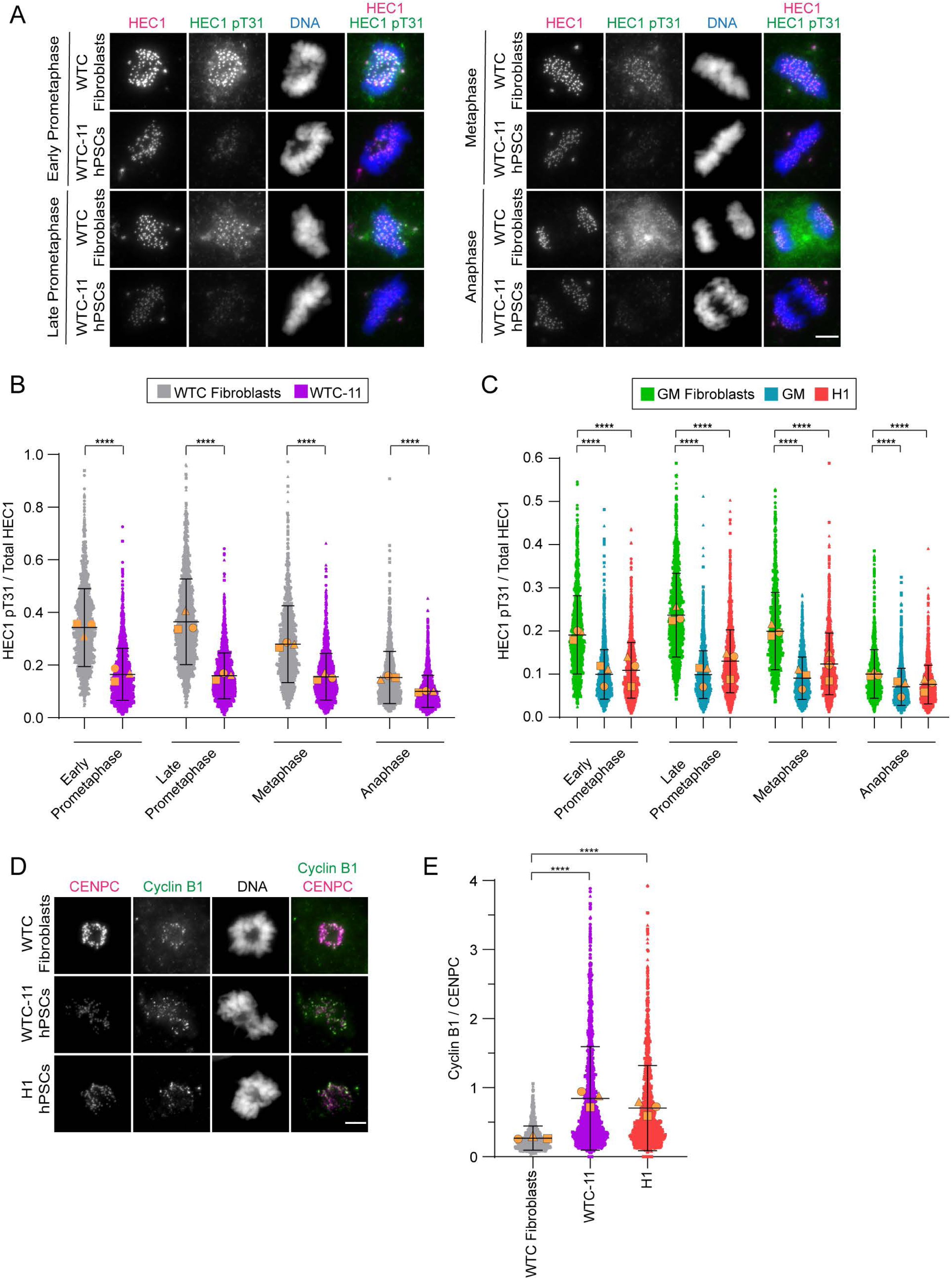
The HEC1 T31 Cyclin B/Cdk1 site is hypophosphorylated at mitotic kinetochores of hPSCs. (**A**) Representative images of mitotic WTC-11 hPSCs and isogenic WTC fibroblasts from early prometaphase to anaphase. Shown is DNA (blue), HEC1 (magenta), and HEC1 phospho-T31 (green). The brightness and contrast for the HEC1 and HEC1 T31 images are scaled equivalently between WTC-11 hPSCs and WTC fibroblasts for comparison. Scale bar 5 μm. (**B**-**C**) Quantification of HEC1 T31 phosphorylation normalized to total HEC1 levels at kinetochores from early prometaphase to anaphase in WTC-11 and WTC fibroblasts (**B**) and GM hPSCs, GM fibroblasts, and H1 hPSCs (**C**). Shown are individual kinetochore intensities and superimposed means of individual replicates (orange shapes). n > 900 total kinetochores from 20 cells per mitotic stage for each replicate with 15-25 kinetochores quantified per cell; mean ± SD. (**D**) Representative images of prometaphase WTC-11 and H1 hPSCs and WTC fibroblasts. Shown is DNA (grayscale), CENP-C (magenta), and Cyclin B1 (green). The brightness and contrast for the CENP-C and Cyclin B1 images are scaled equivalently between hPSCs and fibroblasts for comparison. Scale bar 5 μm. (**E**) Quantification of Cyclin B1 levels normalized to CENP-C levels at prometaphase kinetochores in WTC-11 and H1 hPSCs and WTC fibroblasts. Shown are individual kinetochore intensities and superimposed means of individual replicates (orange shapes). n > 900 total kinetochores from 20 prometaphase cells for each replicate with 15-25 kinetochores quantified per cell; mean ± SD. Three independent biological replicates (**B**-**C** and **E**); ****p < 0.0001 using an unpaired Welch’s t-test (**B**-**C** and **E**).

Both isoforms, Cyclin B1 and Cyclin B2, localize to kinetochores in prometaphase but progressively decline at kinetochores as correct bioriented attachments form (Bentley et al., 2007; Alfonso-Pérez et al., 2019; Liu et al., 2022). Prior results did not distinguish if HEC1 T31 is phosphorylated by Cyclin B1/Cdk1 and/or Cyclin B2/Cdk1 (Kucharski et al., 2022), so we measured prometaphase kinetochore levels of both normalized to CENP-C to compare their abundance in hPSCs vs. somatic cells. In contrast to Aurora B kinase, both Cyclin B1 and Cyclin B2 were significantly enriched at prometaphase kinetochores of hPSCs compared to somatic cells (Figures 3D-E and S4C-D). Cdk1 enzymatic activity requires a Cyclin binding partner (Malumbres, 2014), so these results indicate that Cdk1 activity is not limiting HEC1 T31 phosphorylation in hPSCs. Combined, our findings suggest that decreased HEC1 phosphorylation, including S44 and T31, at kinetochores of hPSCs increases HEC1 binding affinity for k-MTs during mitosis. This in turn results in more stable k-MT attachments which reduces merotelic error correction and causes a high rate of lagging chromosomes for hPSCs.

### Increasing HEC1 phosphorylation in hPSCs improves chromosome segregation fidelity

If a pluripotent state confers a high rate of lagging chromosomes due to HEC1 hypophosphorylation then increasing HEC1 phosphorylation at kinetochores of hPSCs will decrease their rate of lagging chromosomes. To test this prediction, we focused on the HEC1 T31 site because the precise phospho-regulation of T31 is critical for high mitotic fidelity as a single non-phosphorylatable alanine substitution at this site increases lagging chromosome rates (Kucharski et al., 2022). Phosphorylation levels are determined by the net sum of kinase and phosphatase activities. Our results show that Cyclin B/Cdk1 activity at kinetochores of hPSCs is not limiting HEC1 T31 phosphorylation (Figures 3D-E and S4C-D) suggesting that elevated phosphatase activity is responsible for hypophosphorylation at this site.

During mitosis, the phosphatases PP1 and PP2A localize to centromeres/kinetochores and are responsible for dephosphorylating centromere and kinetochore substrates (Saurin, 2018). PP1 progressively increases at centromeres/kinetochores peaking in metaphase (DeLuca et al., 2011). In contrast, PP2A is enriched at centromeres/kinetochores during prometaphase and declines as bioriented attachments form (Foley et al., 2011), similar to the localization pattern of Cyclin B/Cdk1 at kinetochores. Although it was previously demonstrated that repressing PP2A activity by siRNA depletion of the B56 regulatory subunits did not increase HEC1 T31 phosphorylation at mitotic kinetochores (Kucharski et al., 2022), we retested the effect of PP2A inhibition on HEC1 T31 phosphorylation using a different strategy. We did this because the PP2A-B55 holoenzyme, not PP2A-B56, preferentially dephosphorylates Ser and Thr Cdk1 phosphorylation sites with a proline in the +1 position (Kruse et al., 2020). We acutely inhibited PP2A phosphatase activity in hPSCs and somatic cells for 1.5 h using the catalytic inhibitor LB-100 that selectively inhibits PP2A over other phosphatases (D’Arcy et al., 2017). Total mitotic duration in hPSCs is less than 1 h (Deng et al., 2023), so this time point ensures that cells enter mitosis in the presence of the inhibitor. There was a dose-dependent increase in HEC1 T31 phosphorylation levels at prometaphase kinetochores for WTC fibroblasts and WTC-11 and H1 hPSCs (Figures 4A-B). Interestingly, we also observed a dose-dependent increase in HEC1 T31 phosphorylation at metaphase kinetochores for WTC-11 and H1 hPSCs, but not WTC fibroblasts (Figure S5A). Although PP2A is predominantly localized to kinetochores at prometaphase (Foley et al., 2011), this suggests that PP2A dephosphorylates kinetochore substates at metaphase in hPSCs. In support, it was recently demonstrated that the PP2A-B56 holoenzyme is active at metaphase kinetochores of HeLa cells (Allan et al., 2025) raising the possibility that PP2A-B55 is also active at metaphase kinetochores. HEC1 T31 phosphorylation levels increased at kinetochores of hPSCs but did not reach the same level as WTC fibroblasts (Figures 4B and S5A). Despite this, there was a significant decrease in lagging chromosome rates with increasing T31 phosphorylation (Figure 4C). Note, hPSCs treated with 50 μM LB-100 did not progress into anaphase, so we could not measure lagging chromosome rates at this concentration. Also, we attempted to treat hPSCs for longer-term with LB-100, but hPSCs did not survive even at low concentrations. Collectively, our results show that Cyclin B/Cdk1 phosphorylation of HEC1 T31 is counteracted by PP2A mediated dephosphorylation at kinetochores for both somatic cells and hPSCs. Also, we demonstrate that increasing HEC1 T31 phosphorylation at kinetochores of hPSCs improves their chromosome segregation fidelity.

**Figure 4:**
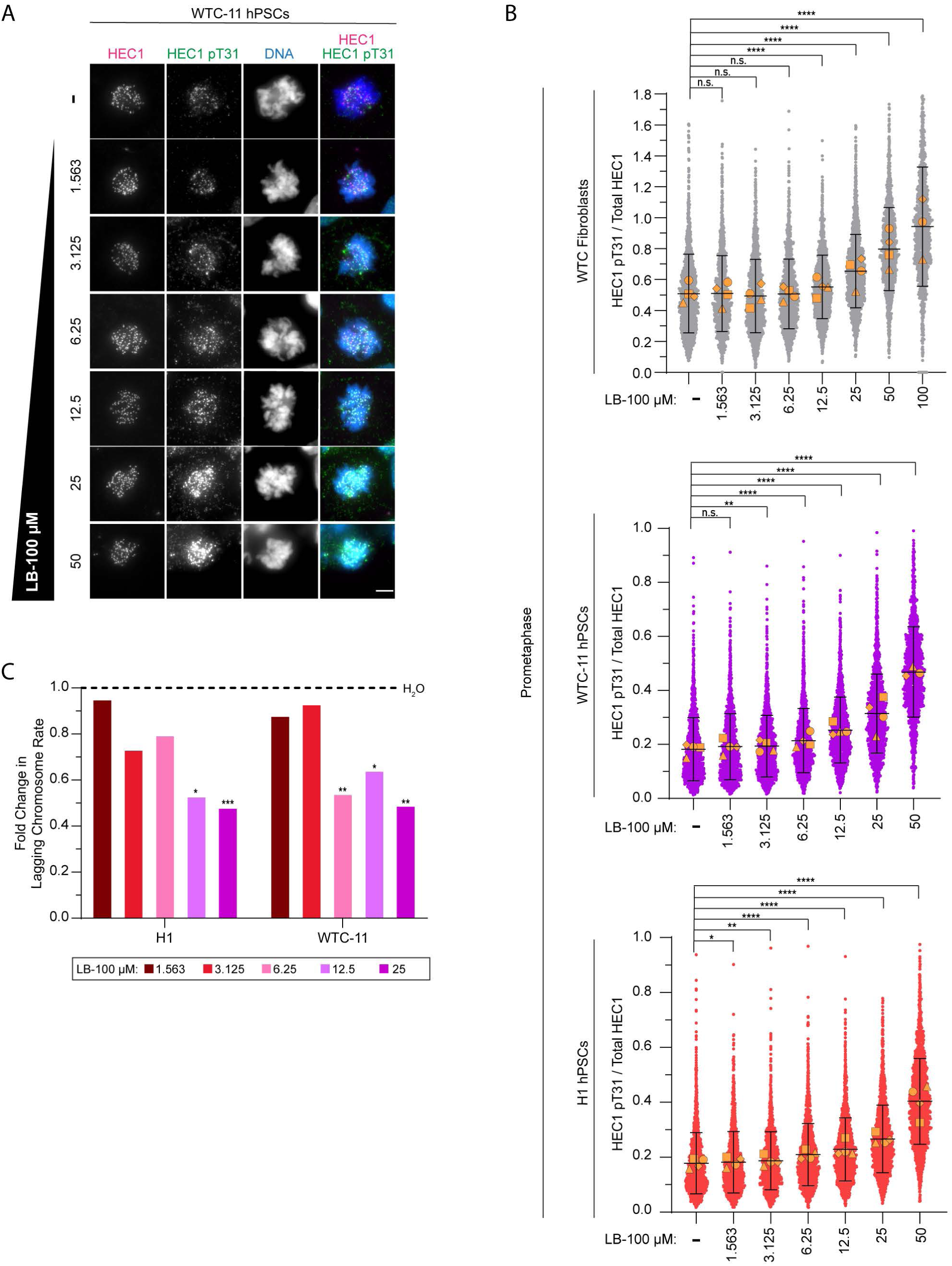
Increasing HEC1 phosphorylation decreases mitotic errors in hPSCs. (**A**) Representative images of control prometaphase WTC-11 hPSCs or cells treated with 2-fold serial dilutions of the PP2A inhibitor LB-100 for 1.5 h. Shown is DNA (blue), HEC1 (magenta), and HEC1 phospho-T31 (green). The brightness and contrast for the HEC1 and HEC1 T31 images are scaled equivalently between WTC-11 hPSCs and WTC fibroblasts for comparison. Scale bar 5 μm. (**B**) Quantification of HEC1 T31 phosphorylation normalized to total HEC1 levels at prometaphase kinetochores of WTC-11 and H1 hPSCs and WTC fibroblasts treated with LB-100. Shown are individual kinetochore intensities and superimposed means of individual replicates (orange shapes). n > 900 total kinetochores from 20 prometaphase cells for each replicate with 15-25 kinetochores quantified per cell; mean ± SD. (**C**) Fold change in the lagging chromosome rate for H1 and WTC-11 hPSCs after 1.5 h treatment with LB-100. For each cell line, the lagging chromosome rate was normalized to an H_2_O control; n > 300 anaphases per condition. Four (**B**) or three independent biological replicates (**C**); n.s. p > 0.05, *p < .05, **p < 0.01, ***p < 0.001, or ****p < 0.0001 using an unpaired Welch’s t-test (**B**) or a two-tailed Fisher’s exact test (**C**).

### Mitotic fidelity depends on the developmental regulation of HEC1 phosphorylation

Our findings above suggest a causal relationship between HEC1 phosphorylation levels and the developmental regulation of chromosome segregation fidelity. If so, then differentiation of hPSCs should increase HEC1 phosphorylation and decrease lagging chromosomes. We previously showed that lagging chromosome rates for hPSCs decreased after 4 days of undirected differentiation using retinoic acid (RA) (Deng et al., 2023), but we did not examine HEC1 phosphorylation levels. Using the same strategy, we measured HEC1 T31 phosphorylation after RA induced differentiation of WTC-11 and H1 hPSCs, which we confirmed resulted in the loss of pluripotency transcription factor expression (Figures S5C-D). HEC1 T31 phosphorylation significantly increased at kinetochores from early prometaphase to anaphase, excluding metaphase, for both WTC-11 and H1 hPSCs upon short-term differentiation (Figures 5A-C and S5B). Crucially, HEC1 T31 phosphorylation increased by ∼20% at prometaphase kinetochores of differentiated hPSCs. This will decrease HEC1 binding affinity for k-MTs resulting in less stable k-MT attachments at prometaphase (Kucharski et al., 2022). This is critical for robust error correction because the frequency of merotelic k-MT attachments is highest in prometaphase (Kabeche and Compton, 2013; Cimini et al., 2003). In agreement, k-MT attachments are least stable in prometaphase for somatic cells that have a low rate of lagging chromosomes (Kabeche and Compton, 2013). Thus, we find that a pluripotent state suppresses HEC1 phosphorylation impairing chromosome segregation fidelity.

**Figure 5.**
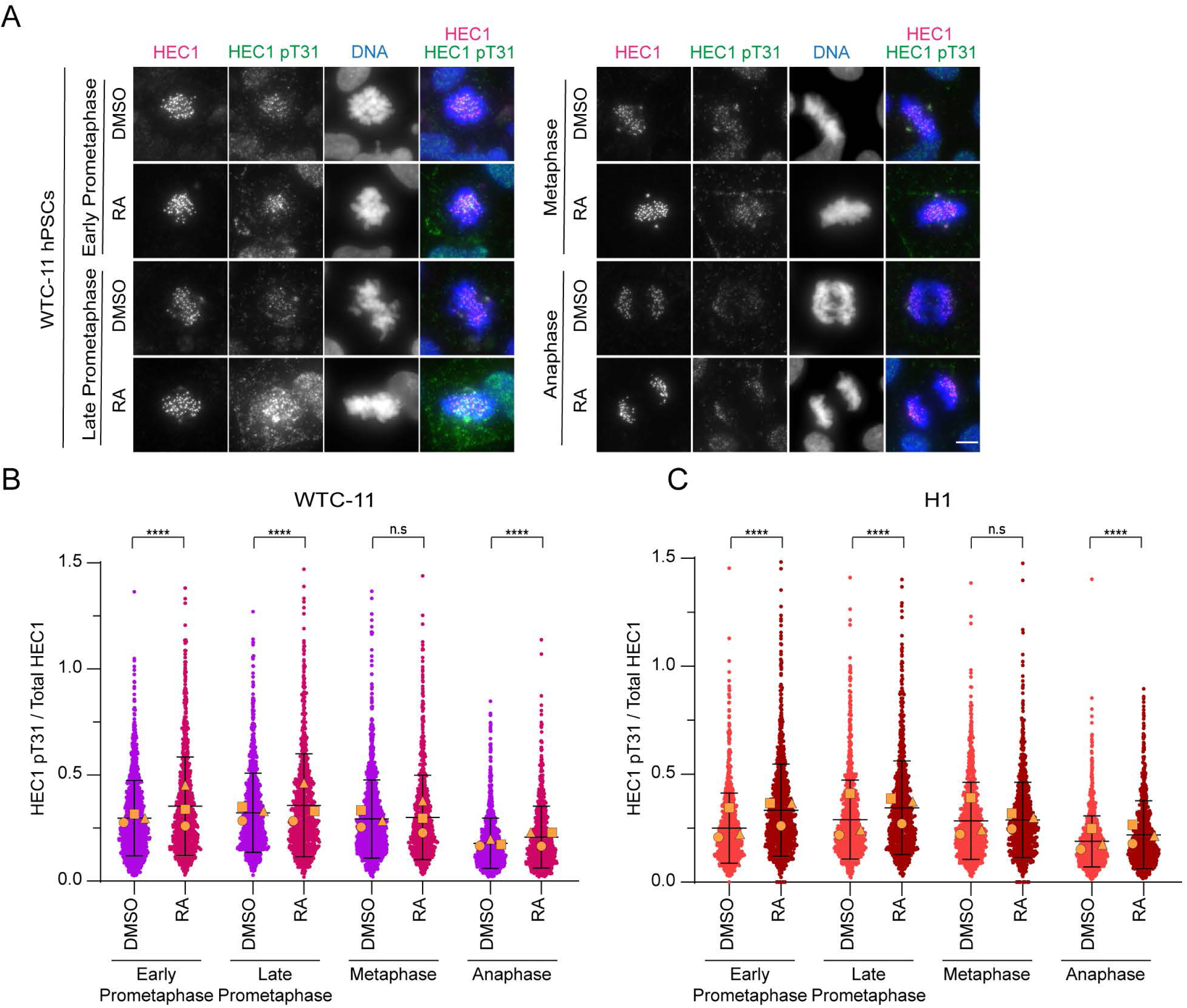
Mitotic HEC1 phosphorylation levels are developmentally regulated. (**A**) Representative images of undifferentiated or differentiated WTC-11 hPSCs from early prometaphase to anaphase after 4-day treatment with DMSO or retinoic acid (RA), respectively. Shown is DNA (blue), HEC1 (magenta), and HEC1 phospho-T31 (green). The brightness and contrast for the HEC1 and HEC1 T31 images are scaled equivalently between undifferentiated and differentiated WTC-11 hPSCs for comparison. Scale bar 5 μm. (**B-C**) Quantification of HEC1 T31 phosphorylation normalized to total HEC1 levels at kinetochores from early prometaphase to anaphase in undifferentiated and differentiated WTC-11 (**B**) and H1 hPSCs (**C**). Shown are individual kinetochore intensities and superimposed means of individual replicates (orange shapes). n > 900 total kinetochores from 20 cells per mitotic stage for each replicate with 15-25 kinetochores quantified per cell; mean ± SD. Three independent biological replicates (**B-C**); n.s. p > 0.05 or ****p < 0.0001 using an unpaired Welch’s t-test (**B-C**).

## Discussion

Faithful chromosome segregation requires both proper centromere and kinetochore formation and phospho-regulation to promote efficient k-MT error correction and establish bi-oriented k-MT attachments. Here, we show that in non-transformed human cells mitotic centromere and kinetochore structural organization and phosphorylation are developmentally regulated (Figure 6). Importantly, genetic variation or artefacts of the reprogramming process cannot explain our results as we compare isogenic human induced pluripotent stem cells and somatic cells and reach similar conclusions in embryonic stem cells. Also, we manipulate developmental state by acutely differentiating hPSCs ruling out the possibility of selection during growth in culture explaining our findings.

**Figure 6:**
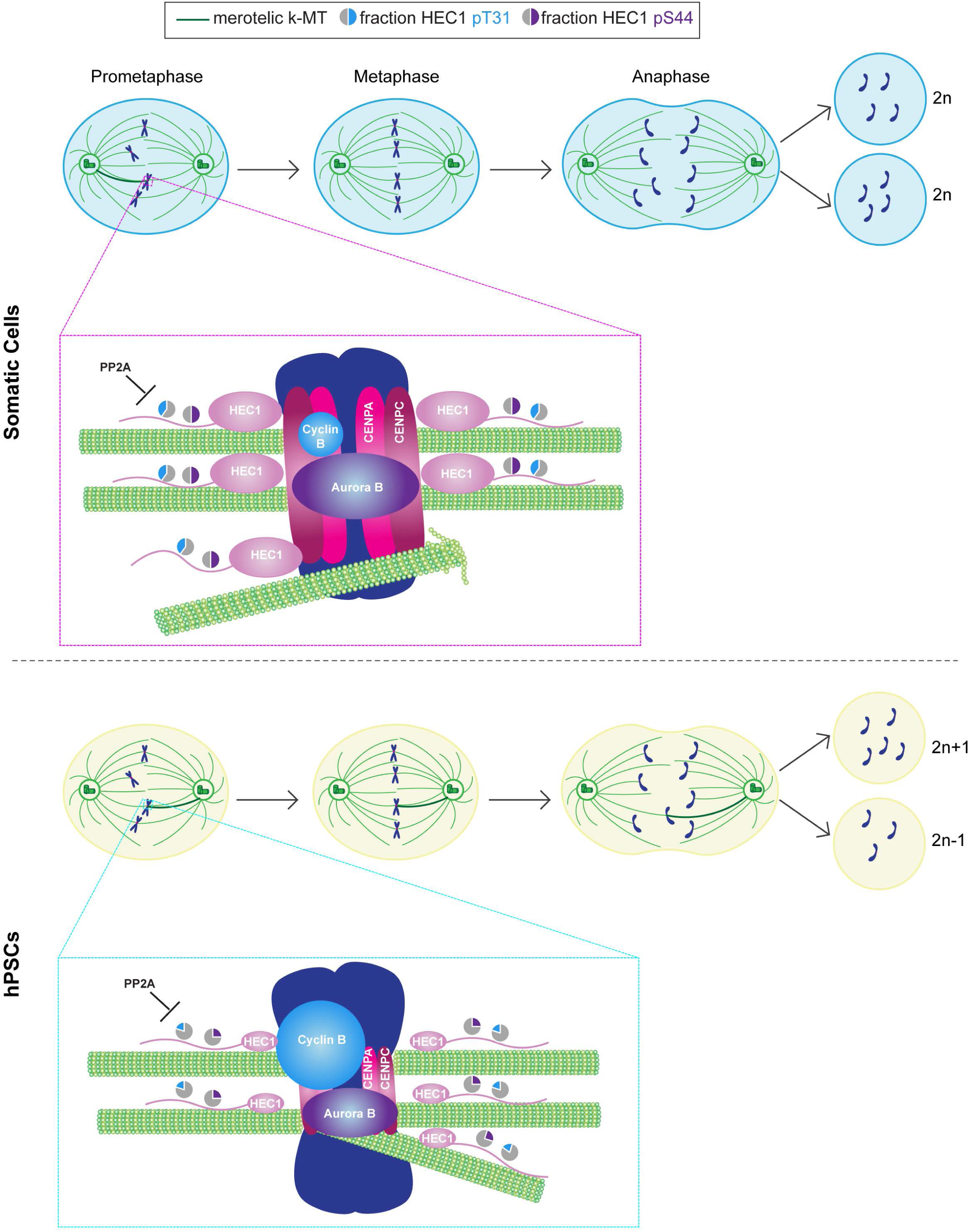
Model for the developmental regulation of mitotic fidelity. Cartoon illustrating the developmental regulation of centromere and kinetochore structural organization and phosphorylation of HEC1. The model compares prometaphase centromeres and kinetochores between somatic cells (upper panel inset) and hPSCs (lower panel inset) when merotelic errors are highest and error correction is most robust to ensure high miotic fidelity. The size of shapes for hPSCs corresponds to relative protein levels compared to somatic cells. Pie charts show the fraction of HEC1 phosphorylated at T31 (blue) or S44 (purple). Although there are differences in the stochiometric abundance of CENP-A, CENP-C, and HEC1 between somatic cells and hPSCs this is not the cause of chromosome segregation errors for hPSCs. Rather, HEC1 is hypophosphorylated at prometaphase kinetochores of hPSCs which increases HEC1 binding affinity for k-MTs causing hyperstable k-MT attachments. Hyperstable k-MT attachments decrease the rate of merotelic error correction by preventing MT depolymerization and detachment. This leads to the persistence of merotelic k-MT attachments (shown as dark green k-MT) and an increase in lagging chromosomes in anaphase for hPSCs. Finally, lagging chromosomes in anaphase can produce aneuploid daughter cells with chromosome gains (2n+1) or losses (2n-1) contributing to hPSC genomic instability.

With respect to centromere and kinetochore structural organization, we show that a pluripotent state causes a decrease in the mitotic abundance of CENP-A centromeric chromatin, centromeric levels of CENP-C, and the outer kinetochore protein HEC1. Also, we demonstrate that CENP-C and HEC1 levels at kinetochores in mitosis scale with the amount of CENP-A chromatin, but it is not a strictly proportional relationship. CENP-A chromatin forms the foundation upon which nearly all other centromere and kinetochore proteins assemble, so likely a pluripotent state leads to a decrease in additional proteins at mitotic centromeres and kinetochores. In support, previous work showed that CENP-T centromere abundance is lower in interphase for hPSCs vs. somatic cells (Milagre et al., 2020).

Interestingly, male undifferentiated mouse germ cells have elevated levels of total nuclear CENP-A in interphase compared to differentiating germ cells and surrounding testicular somatic cells. Also, undifferentiated female mouse germ cells have increased total nuclear CENP-A levels in interphase compared to surrounding somatic cells (Tower et al., 2025). Assuming these findings reflect relative protein differences at mitotic centromeres as well this indicates that there is remarkable plasticity in centromere/kinetochore structural organization with respect to their overall stoichiometry, but the relationship between organization and different developmental states is context dependent. The number of MT attachments positively correlates with kinetochore area (Drpic et al., 2018), so a downstream prediction of these findings is that the number of k-MTs per chromosome required for segregation varies widely among different human cell types. Our data suggests that hPSCs have fewer k-MTs per chromosome compared to somatic cells, but this has not been directly tested.

Why and how developmental programs alter the quantitative composition of centromeres/kinetochores are open questions. A straight-forward explanation for hPSCs is that the epigenetic and transcriptional differences between pluripotent and differentiated states decrease expression of centromere and kinetochore genes limiting protein abundance. Yet evidence shows that soluble pools of centromere and kinetochore proteins are elevated in hPSCs compared to somatic cells (Milagre et al., 2020). This implicates the pathways facilitating centromere and kinetochore assembly, particularly CENP-A deposition, and disassembly as being differentially regulated. More puzzling is what, if any, is the role and consequences for remodeling centromere/kinetochore stoichiometry among different developmental states. Our data demonstrates that developmentally regulated quantitative changes in centromere/kinetochore composition are not a determinant of mitotic chromosome segregation fidelity. In agreement, total whole-cell and nuclear centromere and kinetochore protein levels decline and mitotic errors increase with aging but increasing CENP-A and CENP-C protein expression in cellular aging models does not improve the frequency of mitotic defects either (Sikder et al., 2025). In contrast, chromosome segregation during meiosis is influenced by the quantity of centromere/kinetochore proteins albeit in the context of meiotic drive where chromosomes with increased abundance of centromere/kinetochore proteins are preferentially transmitted to the egg (Akera et al., 2019).

Rather, our results demonstrate that the developmental regulation of centromere and kinetochore function via phosphorylation determines mitotic fidelity. We find that a pluripotent state decreases HEC1 phosphorylation, at least for the T31 and S44 sites, while differentiation and loss of pluripotency increases phosphorylation, at least for T31. We speculate that phosphorylation levels of additional HEC1 N-terminal tail sites at mitotic kinetochores are similarly developmentally regulated. In support, phosphoproteomics analysis of asynchronous populations show that HEC1 S69 phosphorylation increases in RA differentiated hPSCs compared to undifferentiated hPSCs (Brill et al., 2009).

Collectively, the cumulative phospho-occupancy of HEC1 N-terminal tail sites is maintained within a precise, narrow dynamic range to promote mitotic fidelity in somatic cells. There is only a ∼20% difference between binding affinity extremes with no k-MT attachments and proper bioriented k-MT attachments (Kucharski et al., 2022). Assuming a similar dynamic range in phospho-occupancy extends to hPSCs, then the relative decrease in HEC1 phosphorylation levels for hPSCs is within physiologic range to increase k-MT binding affinity. The downstream consequence of hyperstable k-MT attachments is a decrease in merotelic error correction accounting for their high rate of lagging chromosomes. In agreement, our results support a causal relationship between the developmental regulation of HEC1 phosphorylation and chromosome segregation error rates because we find that differentiation of hPSCs increases HEC1 phosphorylation and decreases lagging chromosomes (Deng et al., 2023).

In addition, we provide mechanistic insights into the developmental regulation of HEC1 phosphorylation. HEC1 phosphorylation at mitotic kinetochores is determined by competing kinase and phosphatase networks, and our results indicate that developmental programs impact activities of both. On one hand, a pluripotent state reduces Aurora B at centromeres/kinetochores which is consistent with diminished HEC1 S44 phosphorylation. On the other hand, a pluripotent state increases Cyclin B/Cdk1 at kinetochores indicating that elevated phosphatase activity is primarily responsible for HEC1 T31 hypophoshorylation. In agreement, we find that PP2A inhibition increases HEC1 phosphorylation at prometaphase and metaphase kinetochores of hPSCs, but not metaphase kinetochores of somatic cells, indicative of higher overall PP2A activity in hPSCs. This could be driven by increased PP2A catalytic activity and/or abundance at kinetochores of hPSCs compared to somatic cells, but a lack of validated antibodies for the catalytic subunit prevents us from easily distinguishing these possibilities (Frohner et al., 2020).

We exclusively analyzed HEC1 phospho-T31 levels because dysregulation of phosphorylation at this site alone is sufficient to increase chromosome segregation errors (Kucharski et al., 2022), but PP2A inhibition likely influences additional HEC1 phosphorylation sites. In support, prior work showed that depletion of PP2A-B56 activity elevated HEC1 S44 phosphorylation at kinetochores of somatic and cancer cells (Kucharski et al., 2022). Thus, our results showing a decrease in mitotic errors for hPSCs upon PP2A inhibition likely reflect the effect of a net combined increase in HEC1 phosphorylation. Although we cannot rule out the possibility that PP2A inhibition effects substrates in addition to HEC1 to improve chromosome segregation fidelity in hPSCs, these findings advocate for PP2A inhibition as a strategy to preserve hPSC genome integrity and prevent aneuploidy. A challenge is that catalytic inhibitors will affect non-mitotic PP2A functions as well.

Currently, it is unknown which developmental pathways regulate the mitotic kinase and phosphatase networks that impact HEC1 phosphorylation. Recent evidence connects the WNT, BMP, and FGF developmental signaling pathways to chromosome segregation fidelity of hPSCs via their regulation of DNA replication dynamics in S-phase (Jaime-Soguero et al., 2024). DNA replication stress is proposed to impair mitotic fidelity through transient multipolar intermediates that increase the formation rate of merotelic k-MT attachments (Wilhelm et al., 2019), but it is unknown if these or alternative developmental signaling pathways also influence the correction rate of merotelic attachments through mitotic HEC1 phosphorylation levels. Future studies will address this possibility. Moreover, the regulation of mitotic kinases and phosphatases by developmental signaling pathways likely leads to differential phosphorylation for additional centromere/kinetochore proteins and a more global influence on centromere/kinetochore function. In support, other centromere and kinetochore proteins in addition to HEC1 are differentially phosphorylated between asynchronous populations of differentiated and undifferentiated hPSCs (Brill et al., 2009). In conclusion, we find that the developmental regulation of kinase and phosphatase networks controlling chromosome segregation governs mitotic fidelity for non-transformed human cells.

## Materials and methods

### Cell culture

H1/WA01 human embryonic stem cells are available from WiCell Research Institute. WTC-11 (GM25256) and GM (GM23476) human induced pluripotent stem cells and GM fibroblasts (GM04506) are available from the Coriell Institute for Medical Research. WTC fibroblasts were obtained from the Gladstone Stem Cell Core. We generated H1 and WTC-11 hPSCs that stably overexpress EGFP-CENP-A (H1 CENP-A OE and WTC-11 CENP-A OE) by co-transfecting cells with the PiggyBac EF1α-EGFP-CENP-A and transposase plasmids.

All cell lines were cultured at 37°C in a humidified atmosphere with 5% CO_2_ and routinely validated as mycoplasma free (Sigma-Aldrich Mycoplasma Kit #MP0035). H1, WTC-11, and GM hPSCs were grown on hESC qualified Matrigel (Corning #354277) in mTeSR1 media (StemCell Technologies #85870). Media was changed daily. H1 and WTC-11 hPSCs were passaged with veresene and GM hPSCs were passaged with ReLeSR (Stemcell Technologies #05872). H1 CENP-A OE hPSCs were grown in mTeSR1 media supplemented with 0.3 ug/mL puromycin (Thermofisher Cat# A1113803) and WTC-11 CENP-A OE hPSCs were grown in mTeSR1 media supplemented with 0.5 ug/mL puromycin. For routine passaging and seeding of H1 CENP-A OE and WTC-11 CENP-A OE hPSCs, puromycin was added to mTeSR1 media after 24 h of recovery. Seeding and culturing of H1 CENP-A OE and WTC-11 CENP-A OE hPSCs on coverslips was done in mTeSR1 media without puromycin to remove puromycin as a confounding variable. Media was changed daily and cells were fixed after 48 h. WTC fibroblasts were cultured in DMEM with 10% FBS, 2 mM GlutaMAX-1 (ThermoFisher #35050061), 0.1 mM MEM nonessential amino acids, 100 U/mL penicillin, and 100 μg/mL streptomycin and passaged with TrypLE Select (ThermoFisher #12563011). GM fibroblasts were cultured in EMEM with 15% FBS, 100 U/mL penicillin, and 100 μg/mL streptomycin and passaged with 0.05% Trypsin.

For all-trans retinoic acid (RA) (Sigma #R2625) differentiation, H1 and WTC-11 hPSCs were plated on Matrigel-coated 18 mm glass coverslips and grown in mTeSR1 in 12-well cell culture plates. After 24 h,1 μM all-trans RA was added to fresh mTeSR1 media. Media with 1 µM RA was changed daily for 4 days prior to fixation. For LB-100 treatment, H1 and WTC-11 hPSCs were plated on coverslips as described above. After 24 h, 1.5 µM to 100 µM LB-100 (MedKoo Cat#: 206834) or water was added to fresh mTeSR1 media for 1.5 h prior to fixation.

### Stable cell line construction

EGFP-CENP-A plasmid (gift from Aaron Straight) was linearized and PCR amplified using the following oligonucleotides to add Nhe1 and Not1 restriction sites: EGFP CENP-A-Nhe1 5’-TCA GCA GCT AGC ATG GTG AGC AAG GGC-3’ and EGFP-CENP-A-Not1 5’-TAA TAT GCG GCC GCT TAG CCC AGG CCC TCT T-3’. The resulting Nhe1-EGFP-CENP-A-Not1 fragment was inserted into the multiple cloning site of the PiggyBac vector, pEF1α-MCS-EF1α-Puro (our lab) through restriction digest and ligation. The resulting PiggyBac plasmid, EF1α-EGFP-CENP-A-EF1α-Puro was verified by sequencing.

The PiggyBac EF1α-EGFP-CENP-A-EF1α-Puro plasmid and a PiggyBac transposase plasmid (gift from Aaron McKenna) were introduced by electroporating cells using the LONZA 4D nucleofector with the P3 Primary Cell 4D-Nucleofector® X Kit L (Cat# No V4XP-3012) using the pulse code CB-150. Twenty-four to seventy-two hours after recovery from nucleofection, EGFP-CENP-A expressing cells were selected with puromycin.

### Immunofluorescence (IF)

H1, WTC-11, H1 CENP-A-OE, WTC-11 CENP-A-OE, and GM hPSCs were plated as aggregates on Matrigel-coated 18 mm glass coverslips in 12-well cell culture plates. WTC fibroblasts and GM fibroblasts were plated on standard 18 mm glass coverslips in 12-well cell culture plates. For staining of pluripotency transcription factors (OCT4, NANOG, SOX2, MYC), and mitotic errors, cells were fixed with 3.5% paraformaldehyde for 5 mins at room temperature. Following fixation, cells were permeabilized with TBS + 0.1% Triton X-100 for 2 x 5 min washes. Next, blocking was done for 30 mins at room temperature in TBS + 0.1% Triton X-100 + 2% BSA or TBS + 0.1% Triton X-100 + 2% BSA +10% donkey serum for SOX2 and MYC. Samples were then incubated with primary antibodies in blocking buffer overnight at 4°C. Subsequently, samples were washed in blocking buffer 3 x 5 mins at room temperature and incubated with fluorescent secondary antibodies and DAPI in blocking buffer for 1 h at room temperature. Samples were then washed in blocking buffer 2 x 5 mins, permeabilization buffer 1 x 5 mins, TBS 1 x 5 mins, and mounted on glass slides with ProLong Gold antifade (Thermo Fisher Scientific #P36934). For HEC1, Cyclin B, CENP-A, CENP-C, and Aurora B IF, cells were pre-extracted in PHEM buffer (60 mM Pipes, 25 mM Hepes, pH 6.9, 10 mM EGTA, 4 mM MgSO4, 1% Triton X-100, and 10 mM glycerol 2-phosphate) for 10 min at 4°C, before being fixed with 3.5% PFA for 5 mins at room temperature. Once fixed, IF was performed as described above.

### Antibodies

The following antibodies were used for IF: rabbit anti-OCT4 (Abcam #ab19857; 5 μg/mL), mouse anti-NANOG (Abcam #ab173368; 1:200), rabbit anti-MYC (Cell Signaling #13987; 1:1600), mouse anti-SOX2 (R&D Systems #MAB2018; 1:50), mouse anti-CENP-A (Thermo Scientific #MA1-20832; 1:400), guinea pig anti-CENP-C (MBL #PD030; 1:100), chicken anti-GFP (Abcam #ab13970; 1:1000), rabbit anti-Aurora B (Novus #NB100-294, 1:1000), mouse anti-Cyclin B1 (Santa Cruz #sc-245; 1:200), rabbit anti-Cyclin B2 (Abcam #ab185622; 1:500), rat anti-tyrosinated tubulin (Santa Cruz #sc-53029; 1:1000 or Novus #NB600-506; 1:2500), mouse anti-HEC1 (Santa Cruz #sc-515550; 1:500), rabbit anti-HEC1 S44 (gift of Jennifer DeLuca; 1:500), and rabbit anti-HEC1 pT31 (Duane Compton; 1:100,000).

For secondary antibodies, all dilutions were at 1:1000: donkey anti-mouse Alexa Fluor 488 (ThermoFisher #A-21202), goat anti-rabbit Alexa Flour 488 (ThermoFisher #A-32731), goat anti-chicken Alexa Fluor 488 (ThermoFisher #A-32931), goat anti-mouse 594 Alexa Fluor (ThermoFisher #A-32742), goat anti-rabbit Alexa Fluor 594 (ThermoFisher #A-11037), goat anti-rat Alexa Fluor 594 (ThermoFisher #A-11007), goat anti-guinea pig Alexa Fluor 594 (ThermoFisher #A-11076), donkey anti-mouse Alexa Fluor 647 (ThermoFisher #A-31571), donkey anti-rabbit Alexa Fluor 647 (ThermoFisher #A-31573), goat anti-rat Alexa Fluor 647 (ThermoFisher #A-21247), and goat anti-guinea pig Alexa Fluor 647 (ThermoFisher #A-21450).

### Microscopy

All images were acquired using a Nikon Eclipse Ti inverted microscope equipped with an ORCA-Fusion Gen III sCMOS camera (Hamamatsu) and a SOLA LED light engine (Lumencor) controlled by Nikon NIS-Elements software version 5.21.01. Images series were acquired with 0.2 µm Z-steps using a Plan Apo VC 60x, 1.4 NA oil immersion objective (Nikon).

### Image Analysis

Quantifications of protein and protein phosphorylation levels at individual centromeres and kinetochores, with the exception of Aurora B, were measured using the Centromere Recognition and Quantification (CRaQ) Fiji plugin (Bodor et al., 2012). In brief, areas bounding centromeres and kinetochores were identified as regions of interest (ROIs) using the CENP-A, CENP-C, or HEC1 signals from maximum intensity z-stack projections. ROIs generated by the plugin were visually confirmed to belong to mitotic cells and not neighboring interphase cells. Background corrected intensity measurements were then calculated as the difference between maximum and minimum intensity values within ROIs. Phosphorylated HEC1 intensity was normalized to total HEC1 intensity or Cyclin B intensity was normalized to CENP-C intensity.

For Aurora B quantification, tubulin staining was used to identify metaphase sister kinetochore pairs through individual z-stack. A circular region 5 pixels in diameter was manually drawn around CENP-C foci of sister kinetochore pairs. A FIJI macro was created to draw a 2 µm line intersecting both sister foci with the midpoint of the line corresponding to the midpoint between the sister foci. A line scan was then performed to measure both Aurora B intensity between the CENP-C sister foci and the CENP-C intensity at each foci. The ratio of peak Aurora B intensity over the line scan was then normalized to the CENP-C intensity from the sister foci with the highest value.

For interkinetochore distance (IKD) measurements, tubulin staining was used to identify metaphase sister kinetochore pairs through individual z-stacks. A circular region 5 pixels in diameter was manually drawn around HEC1 foci of sister kinetochore pairs. The centroid X and Y coordinates of each sister was used in the following equation to calculate IKD: SQRT((KineX1-KineX2)^2^+(KineY1-KineY2)^2^).

For spindle size measurement, a circular ROI was drawn around metaphase mitotic spindles encompassing both spindle poles from maximum intensity z-stack projections. The diameter of the spindle was calculated using the Minimum Feret’s Diameter measurement option in FIJI.

Nuclear protein levels of pluripotency transcription factors OCT4, NANOG, SOX2, and MYC and mitotic error quantifications were done as we previously described (Deng et al., 2023).

### Statistical analysis

All statistical tests were performed using Graphpad Prism version 10.6.1. All experiments were repeated for a minimum of three independent biological replicates. At least 20 cells were analyzed per replicate with 15-25 individual kinetochores measured for each cell or at least 10 cells were analyzed per replicate with 5-10 sister kinetochore pairs measured for each cell. For pluripotency transcription factor quantifications, 100 cells per replicate were randomly selected using DAPI staining. For anaphase error rates, 100 anaphases per condition were randomly assessed per replicate. The mean value for individual replicates is superimposed on data plots using orange shapes except for mitotic error quantifications. Statistical significance was calculated using unpaired Welch’s t-test, unpaired one-way ANOVA with Holm-Sadak’s multiple comparison test, or Fisher’s exact test as indicated in figure legends. No outliers were excluded.

### Online supplemental material

Figure S1 (in support of Figure 1) shows that mitotic centromere and kinetochore protein abundance is low for hPSCs compared to isogenic somatic cells and that CENP-C levels scale with CENP-A chromatin. Figure S2 (in support of Figure 1) shows that GFP-CENP-A expressing hPSCs also express pluripotency transcription factors. Figure S3 (in support of Figures 1 and 2) shows that mitotic HEC1 abundance is low for hPSCs compared to isogenic somatic cells and that interkinetochore distance and metaphase spindle length are shorter for hPSCs compared to somatic cells. Figure S4 (in support of Figures 1 and 3) shows that the HEC1 T31 Cyclin B/Cdk1 site is hypophosphorylated at mitotic kinetochores of hPSCs and that Cyclin B2 is increased at prometaphase kinetochores of hPSCs. Figure S5 (in support of Figures 4 and 5) shows that PP2A inhibition increases HEC1 T31 phosphorylation at metaphase kinetochores of hPSCs and that differentiation of hPSCs increases HEC1 T31 phosphorylation at mitotic kinetochores.

## Data availability

Materials and data are available from corresponding author upon request.

## Acknowledgements

We thank Jennifer DeLuca for sharing the HEC1 S44 phospho-antibody, Yina Huang for sharing equipment, and Aaron Straight and Aaron McKenna for sharing plasmids. Also, we thank Godek lab members, Compton lab members, Amanda Amodeo, and Aaron McKenna for their helpful discussions. This work was supported by the Genomics and Molecular Biology Shared Resources facilities at the Dartmouth Cancer Center with NCI Cancer Center Support Grant 5P30 CA023108-41 and the National Institute of Health R01HD101436 to KMG.

## Author contributions

Conceptualization-KMG; Methodology-KMG, BGS; Validation-KMG, BGS; Formal Analysis-BGS; Investigation-BGS; Resources-KMG, DAC; Writing-Original Draft-KMG, BGS; Writing-Review and Editing-KMG, BGS, DAC; Visualization-KMG, BGS; Supervision-KMG; Funding Acquisition-KMG.

## Figure Legends

**Figure S1:**
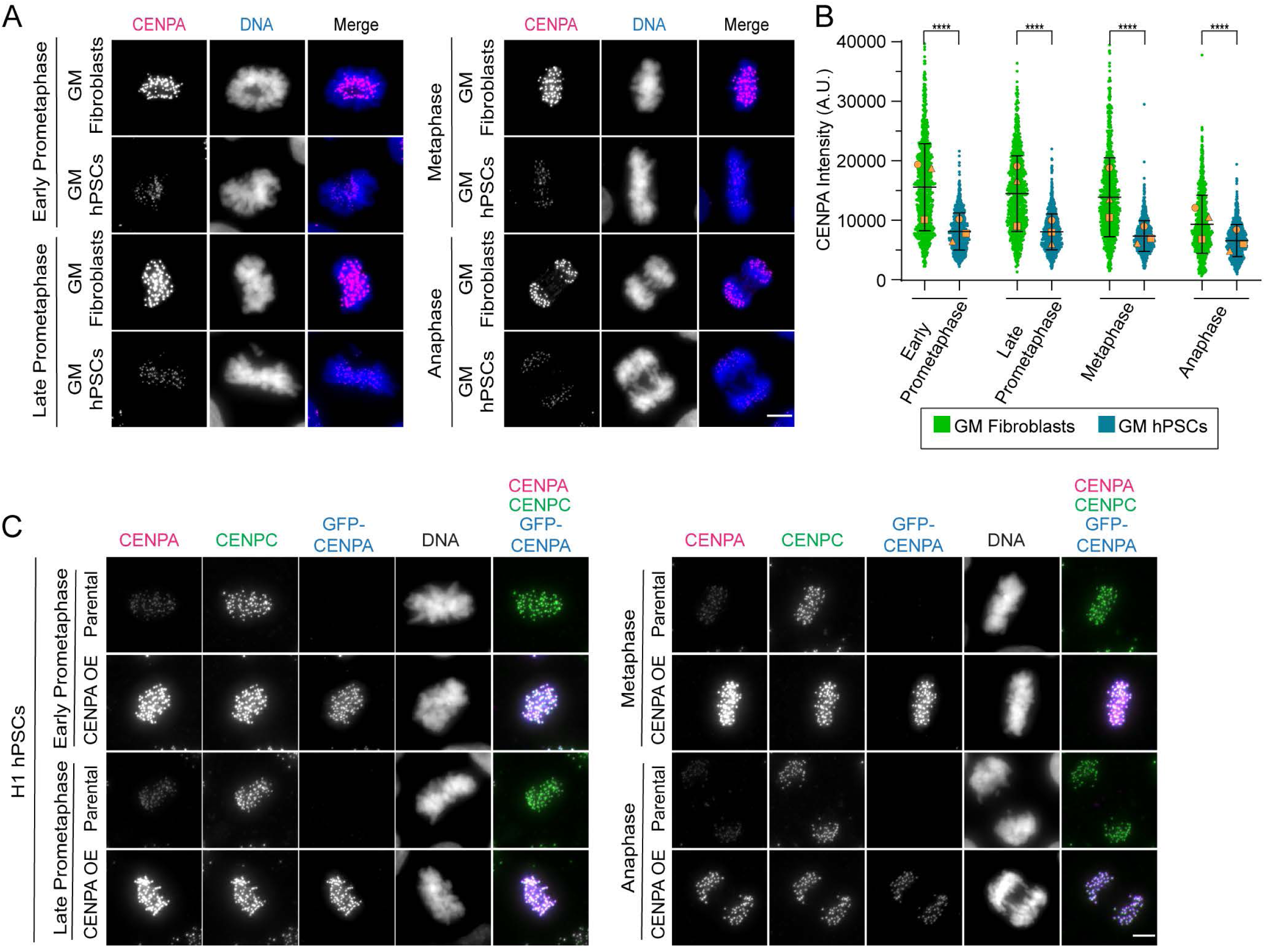
Mitotic centromere and kinetochore protein abundance is low for hPSCs compared to isogenic somatic cells; CENP-C levels scale with CENP-A chromatin. (**A**) Representative images of mitotic GM hPSCs and GM fibroblasts from early prometaphase to anaphase. Shown is DNA (blue) and CENP-A (magenta). The brightness and contrast for the CENP-A images are scaled equivalently between GM hPSCs and GM fibroblasts for comparison. Scale bar 5 μm. (**B**) Quantification of total CENP-A kinetochore levels from early prometaphase to anaphase in GM hPSCs and GM fibroblasts. Shown are individual kinetochore intensities and superimposed means of individual replicates (orange shapes). n > 900 total kinetochores from 20 cells per mitotic stage for each replicate with 15-25 kinetochores quantified per cell; mean ± SD; three independent biological replicates; ****p < 0.0001 using an unpaired Welch’s t-test. (**C**) Representative images of mitotic H1 hPSCs and H1 CENP-A-OE hPSCs from early prometaphase to anaphase. Shown is DNA (grayscale), CENP-A (magenta), CENP-C (green), and GFP (blue). The brightness and contrast for the CENP-A and CENP-C images are scaled equivalently between parental and CENP-A OE hPSCs for comparison. Scale bar 5 μm.

**Figure S2:**
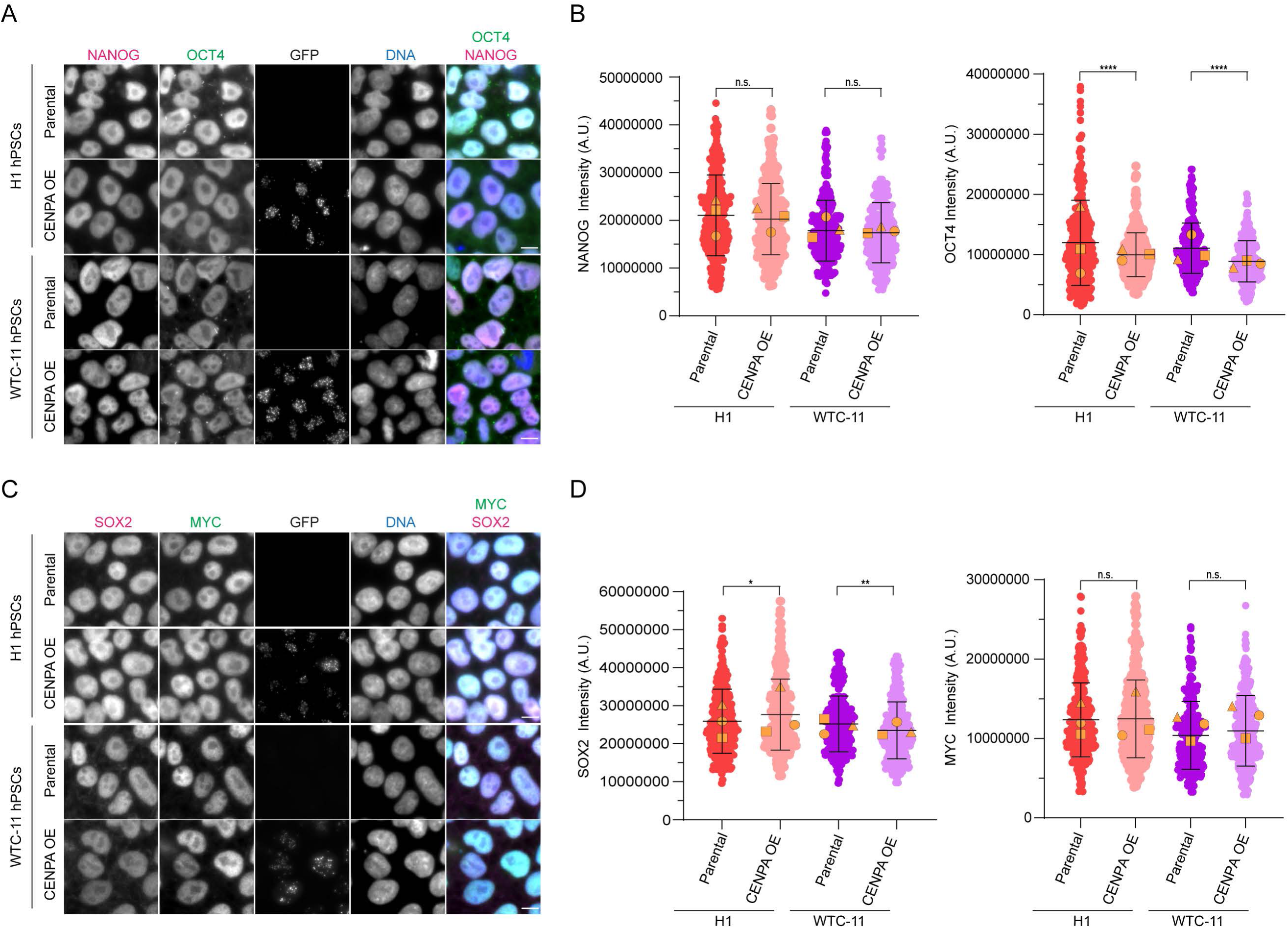
GFP-CENP-A expressing hPSCs also express pluripotency transcription factors. (**A**) Representative images of parental and GFP-CENP-A OE H1 and WTC-11 hPSCs. Shown is DNA (blue), NANOG (magenta), OCT4 (green), and GFP (grayscale). The brightness and contrast for the OCT4 and NANOG images are scaled equivalently between parental and CENP-A OE hPSCs for comparison. Scale bar 10 μm. (**B**) Quantification of nuclear NANOG (left panel) and OCT4 (right panel) levels for parental and GFP-CENP-A OE H1 and WTC-11 hPSCs. Shown are individual nucleus intensities and superimposed means of individual replicates (orange shapes). n = 300 nuclei; mean ± SD. (**C**) Representative images of parental and GFP-CENP-A OE H1 and WTC-11 hPSCs. Shown is DNA (blue), SOX2 (magenta), MYC (green), and GFP (grayscale). The brightness and contrast for the SOX2 and MYC images are scaled equivalently between parental and CENP-A OE hPSCs for comparison. Scale bar 10 μm. (**B**) Quantification of nuclear SOX2 (left panel) and MYC (right panel) levels for parental and GFP-CENP-A OE H1 and WTC-11 hPSCs. Shown are individual nucleus intensities and superimposed means of individual replicates (orange shapes). n = 300 nuclei; mean ± SD. Three independent biological replicates (**B** and **D**); n.s. p > 0.05, *p < .05, **p < 0.01, or ****p < 0.0001 using an unpaired Welch’s t-test.

**Figure S3:**
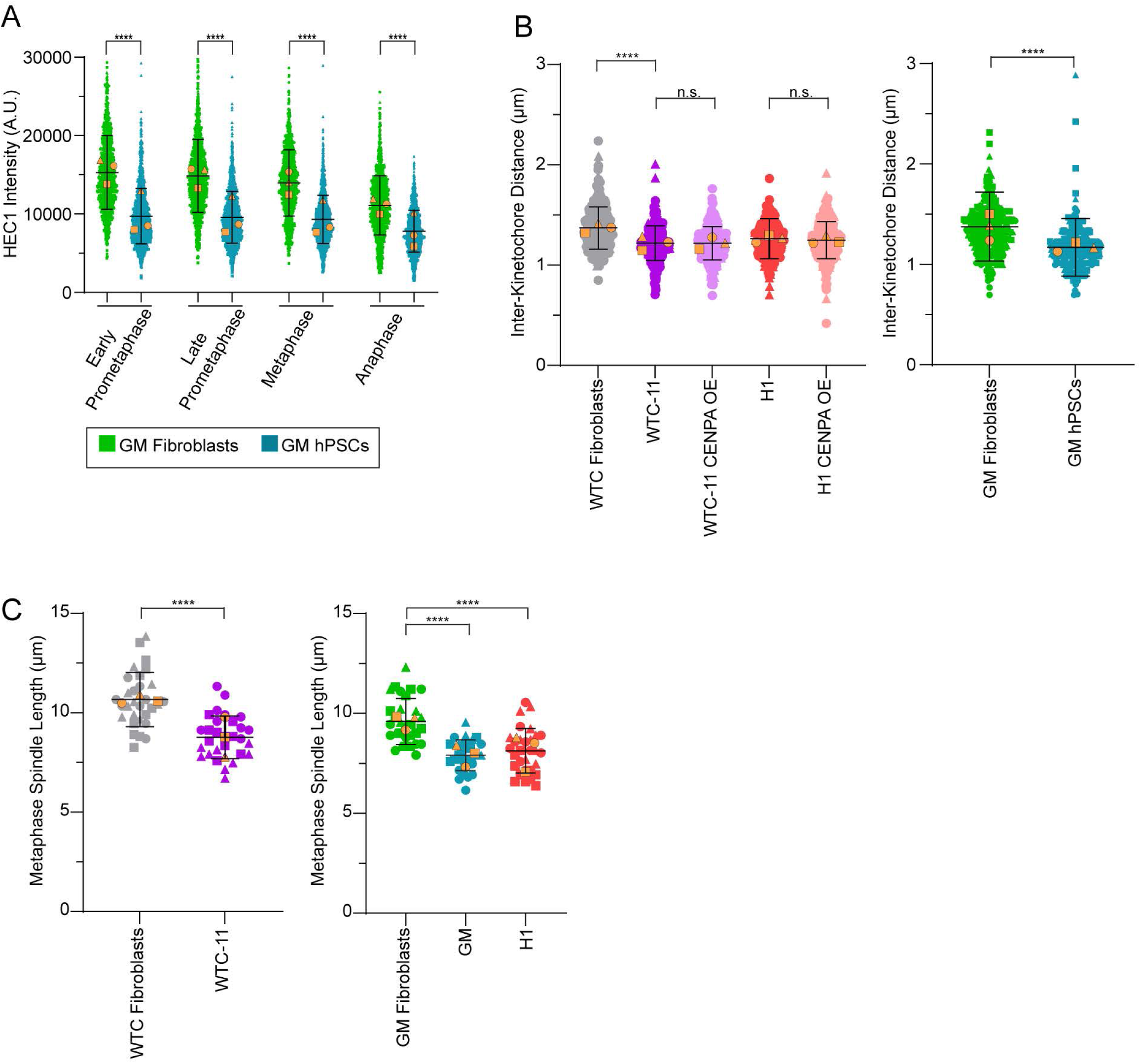
Mitotic HEC1 abundance is low for hPSCs compared to isogenic somatic cells; interkinetochore distance and metaphase spindle length are shorter for hPSCs compared to somatic cells. (**A**) Quantification of total HEC1 kinetochore levels from early prometaphase to anaphase in GM hPSCs and GM fibroblasts. Shown are individual kinetochore intensities and superimposed means of individual replicates (orange shapes). n > 900 total kinetochores from 20 cells per mitotic stage for each replicate with 15-25 kinetochores quantified per cell; mean ± SD. (**B**) HEC1 interkinetochore distance was measured between pairs of sister chromatids at metaphase for H1, H1 CENP-A OE, WTC-11, WTC-11 CENP-A OE, WTC fibroblasts, GM hPSCs, and GM fibroblasts. Shown are sister pairs and superimposed means of individual replicates (orange shapes). n = 300 total sister kinetochore pairs from 10 cells for each replicate with 10 pairs quantified per cell; mean ± SD. (**C**) Metaphase spindle length measured in WTC-11, GM, and H1 hPSCs and WTC fibroblasts and GM fibroblasts. Shown are individual spindles (circles) and superimposed means of individual replicates (orange shapes). n = 30 metaphase spindles from 10 cells for each replicate; mean ± SD. Three independent biological replicates (**B-C**); n.s. p > 0.05 or ****p < 0.0001 using an unpaired Welch’s t-test (**A, B** right plot, and **C**) or using a one-way ANOVA with Holm-Sadak’s multiple comparison test (**B** left plot).

**Figure S4:**
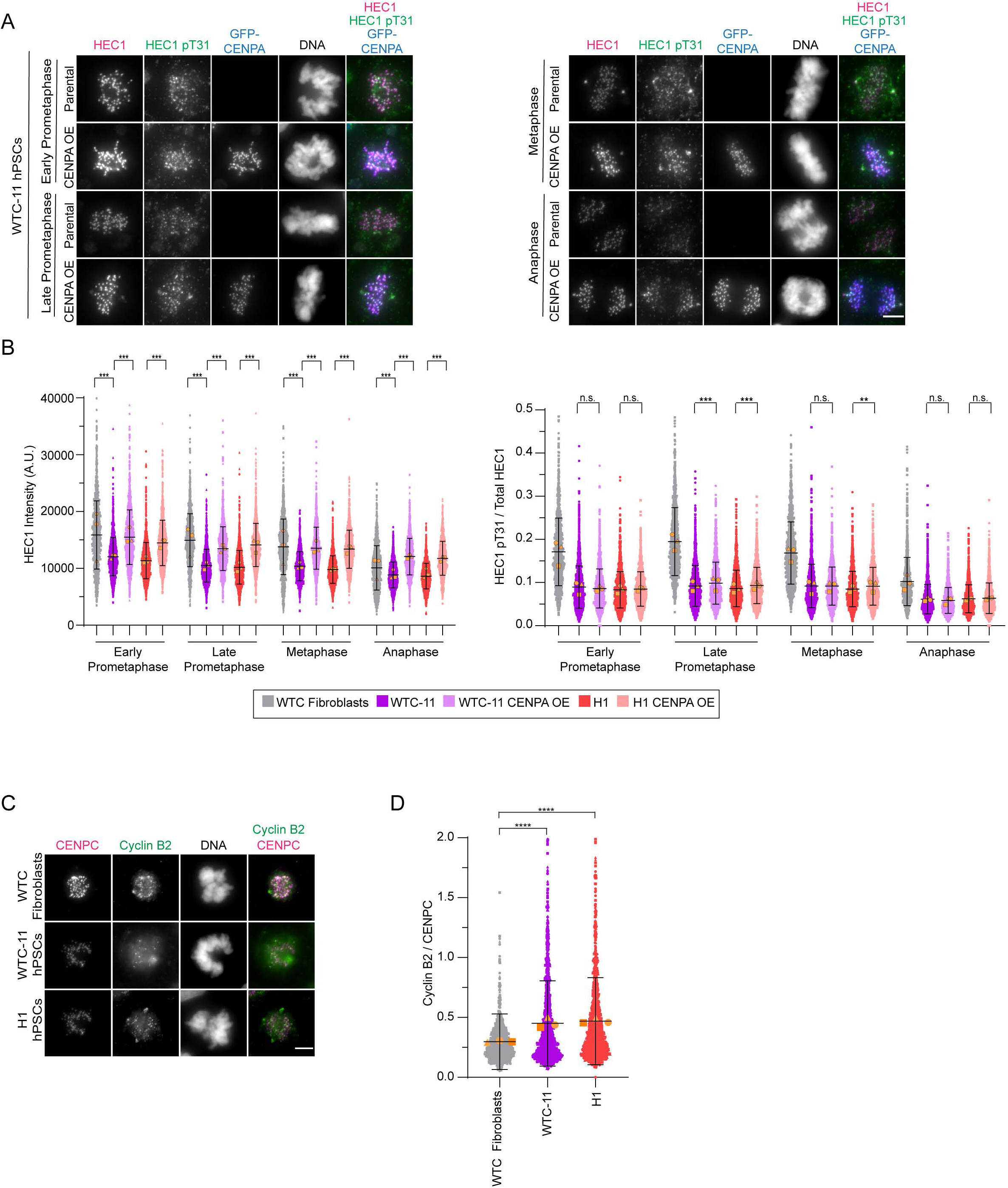
The HEC1 T31 Cyclin B/Cdk1 site is hypophosphorylated at mitotic kinetochores of hPSCs; Cyclin B2 is increased at prometaphase kinetochores of hPSCs. (**A**) Representative images of mitotic WTC-11 and WTC-11 GFP-CENP-A OE hPSCs from early prometaphase to anaphase. Shown is DNA (blue), HEC1 (magenta), and HEC1 phospho-T31 (green). The brightness and contrast for HEC1 and HEC1 T31 images are scaled equivalently between WTC-11 hPSCs and WTC fibroblasts for comparison. Scale bar 5 μm. (**B**) Quantification of HEC1 T31 phosphorylation normalized to total HEC1 levels at kinetochores from early prometaphase to anaphase in WTC-11, WTC-11 GFP-CENP-A OE, H1, and H1 GFP-CENP-A OE hPSCs and WTC fibroblasts. Shown are individual kinetochore intensities and superimposed means of individual replicates (orange shapes). n > 900 total kinetochores from 20 cells per mitotic stage for each replicate with 15-25 kinetochores quantified per cell; mean ± SD. (**C**) Representative images of prometaphase WTC-11 and H1 hPSCs and WTC fibroblasts. Shown is DNA (grayscale), CENP-C (magenta), and Cyclin B2 (green). The brightness and contrast for CENP-C and Cyclin B2 images are scaled equivalently between hPSCs and fibroblasts for comparison. Scale bar 5 μm. (**D**) Quantification of Cyclin B2 levels normalized to CENP-C levels at prometaphase kinetochores in WTC-11 and H1 hPSCs and WTC fibroblasts. Shown are individual kinetochore intensities and superimposed means of individual replicates (orange shapes). n > 900 total kinetochores from 20 prometaphase cells for each replicate with 15-25 kinetochores quantified per cell; mean ± SD. Three independent biological replicates (**B** and **D**); n.s. p > 0.05, **p < 0.01, ***p < 0.001, or ****p < 0.0001 using a one-way ANOVA with Holm-Sadak’s multiple comparison test (**B**) or an unpaired Welch’s t-test (**D**).

**Figure S5:**
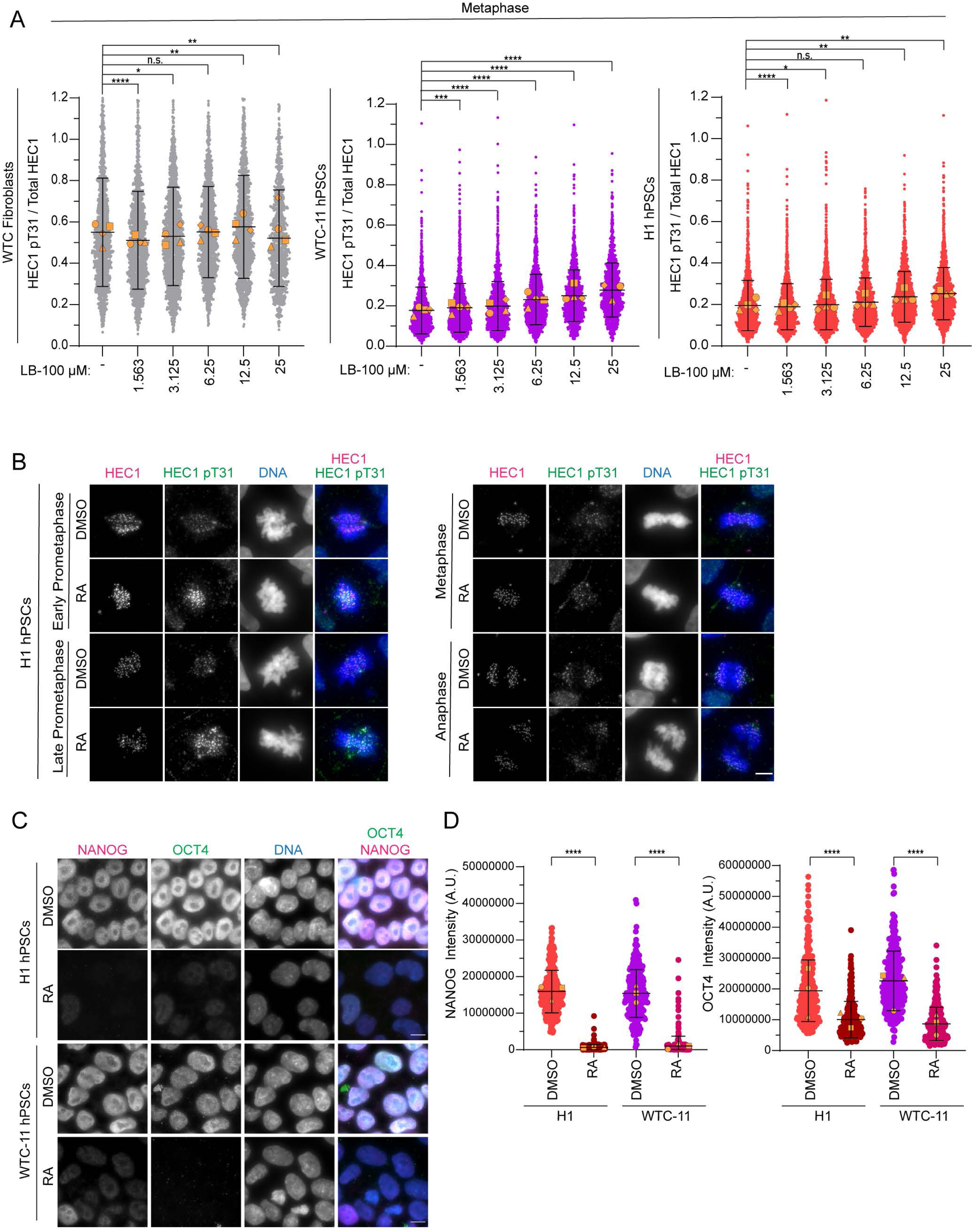
PP2A inhibition increases HEC1 T31 phosphorylation at metaphase kinetochores of hPSCs; differentiation of hPSCs increases HEC1 T31 phosphorylation at mitotic kinetochores. (**A**) Quantification of HEC1 T31 phosphorylation normalized to total HEC1 levels at metaphase kinetochores of untreated WTC-11 and H1 hPSCs and WTC fibroblasts or cells treated with increasing concentrations of the PP2A inhibitor LB-100 for 1.5 h. Shown are individual kinetochore intensities and superimposed means of individual replicates (orange shapes). n > 900 total kinetochores from 20 prometaphase cells for each replicate with 15-25 kinetochores quantified per cell; mean ± SD. (**B**) Representative images of undifferentiated or differentiated H1 hPSCs from early prometaphase to anaphase after 4-day treatment with DMSO or retinoic acid (RA), respectively. Shown is DNA (blue), HEC1 (magenta), and HEC1 phospho-T31 (green). The brightness and contrast for HEC1 and HEC1 T31 images are scaled equivalently between undifferentiated and differentiated H1 hPSCs for comparison. Scale bar 5 μm. (**C**) Representative images of undifferentiated or differentiated H1 and WTC-11 hPSCs after 4-day treatment with DMSO or retinoic acid (RA), respectively. Shown is DNA (blue), NANOG (magenta), and OCT4 (green). The brightness and contrast for OCT4 and NANOG images are scaled equivalently between undifferentiated and differentiated H1 or WTC-11 hPSCs for comparison. Scale bar 10 μm. (D) Quantification of nuclear NANOG (left panel) and OCT4 (right panel) levels for undifferentiated (DMSO) or differentiated (RA) H1 and WTC-11 hPSCs. Shown are individual nucleus intensities (circles) and superimposed means of individual replicates (orange shapes). n = 300 nuclei; mean ± SD. Four (**A**) or three independent biological replicates (**D**); n.s. p > 0.05, *p < .05, **p < 0.01, ***p < 0.001, or ****p < 0.0001 using an unpaired Welch’s t-test (**B** and **D**).

